# Viral diversity and phloem transcriptional changes in Grapevine Shiraz disease infected vines

**DOI:** 10.1101/2024.03.18.585633

**Authors:** Cristobal Onetto, Dilrukshi S. K. Nagahatenna, Yeniu Mickey Wang, Vinay Pagay, Anthony Borneman

## Abstract

Shiraz disease (SD) is a highly destructive disease of grapevines that is associated with Grapevine Virus A (GVA) infection of vineyards in Australia and South Africa. However, little is known about the transcriptional modifications in grapevine phloem tissues induced by SD. In this study, we explored the viral diversity and transcriptional changes linked to SD. Vines symptomatic for SD exhibited higher viral abundance and were also shown to be co-infected with both GVA and grapevine leafroll-associated virus (GLRaV-4) strain 9. Differential gene (DE) expression analysis revealed physiological responses of *Vitis vinifer*a to the infection. Similar to other plant pathogen infections, SD upregulated genes associated with the systemic acquired resistance (SAR) mechanism and downregulated genes related to vine immunity. Additionally, upregulated genes suggests that callose deposition and the blocking of phloem sieve elements are likely employed by *V. vinifera* as a defence strategy to limit the internal spread of SD viruses.

Grapevine Shiraz disease, Grapevine Virus A, Grapevine Leafroll Associated virus, Systemic Acquired Resistance, Transcriptomics

## Introduction

Grapevine (*Vitis vinifera* L.) is a major perennial horticultural crop grown in diverse climatic regions around the world. As grapevine is vegetatively propagated, it is highly susceptible to graft- and vector-transmitted viruses. Currently, 86 phloem-limited viruses have been reported in grapevine, many of which cause economically relevant diseases (Fuchs, 2020).

Shiraz disease (SD) is a highly destructive viral disease of *V. vinifera*, that has so far been reported to occur only in South Africa and Australia (Wu et al., 2020). The grapevine cultivars Shiraz, Merlot, Gamay, Malbec and Viognier are highly susceptible to SD (Goszczynski et al., 2008), with infected vines displaying primary bud necrosis, retarded shoot growth and delayed senescence of crimson-coloured leaves (Wu et al., 2020). However, other cultivars such as Cabernet Sauvignon do not exhibit primary SD-symptoms upon infection (Goszczynski, 2007). While the phenotypic symptoms of the disease have been extensively described, the molecular plant responses to infection remain poorly understood.

The *Vitivirus*, Grapevine Virus A (GVA), belonging to the family *Betaflexiviridae* (Martelli, 1997), is considered the primary causal agent of Shiraz Disease (SD) (Goszczynski and Habili, 2012). Previous studies have shown that SD symptoms are prominent in susceptible cultivars infected with GVA alone or co-infected with variants of Grapevine Leafroll-associated Virus (GLRaV), including GLRaV-1, GLRaV-3, or GLRaV-4 strain 9 (Goszczynski and Habili, 2012; Wu et al., 2020; Wu et al., 2023). Despite this, grapevine leafroll disease, primarily associated with GLRaV-3 variants, presents distinct symptoms from SD (Maree et al., 2013). Some studies suggest that the co-existence of GVA and GLRaV may influence SD pathogenesis (Wu et al., 2020); however, this has not been conclusively demonstrated, and GVA remains the key pathogen linked to SD. Both GVA and GLRaV are phloem-limited viruses transmitted primarily through grafting or insect vectors, such as mealybugs and scale insects (Rosciglione et al., 1983; Martelli, 2001).

GVA has been shown to possess extensive genetic heterogeneity (Goszczynski and Jooste, 2002; Goszczynski et al., 2008; Goszczynski, 2014; Wu et al., 2020), with genetic variants classified into three molecular groups based on nucleotide similarity (Goszczynski and Jooste, 2003). Variants from molecular groups I and III are often asymptomatic whereas variants of group II are found to be closely associated with SD and often symptomatic (Goszczynski and Jooste, 2003; Goszczynski and Habili, 2012; Goszczynski, 2014; Wu et al., 2020).

The molecular responses to grapevine viruses have been minimally investigated, with only a handful of studies examining the alterations in gene expression caused by GLRaV infection (Song et al., 2021). These studies provided insights into the biological functions affected by the disease, including leaf-specific (Gutha et al., 2010) and berry-specific (Ghaffari et al., 2020; Vega et al., 2011) transcriptional changes, which helped to elucidate the genes involved in the symptomology of grapevine leafroll disease. Although SD is a prevalent virus-induced condition in Australian vineyards, no prior studies have explored the molecular stress responses associated with this disease. High-throughput transcriptome sequencing was therefore employed to investigate the SD-induced transcriptomic changes in grapevine phloem tissues, with the goal of gaining an initial understanding of the grapevine’s molecular stress responses to this disease.

## Materials and methods

### 1. Experimental site

The study was conducted over two growing seasons (2020/2021 and 2021/2022) in a commercial vineyard located in Riverland, South Australia. *Vitis vinifera* L. cv. Shiraz vines included in the study were grafted onto K51-40 rootstock and planted in 1998 with a spacing of 3.5 m and 3.5 m within and between rows, respectively, and oriented E-W direction. The vines were drip-irrigated and fertilised according to the standard practices for the region. Vines were spur pruned and shoots produced from bilateral cordons were trained to vertical shoot positioning.

### 2. Identification of viral diversity in asymptomatic and Shiraz diseased vines

Over the course of both growing seasons, all vines were visually assessed for known SD symptoms (Qi et al. 2020), including retarded shoot growth, partial lignification patterns in canes, and retention of crimson-coloured leaves in winter. Depending on the severity of the symptoms at the harvest stage (in March), cane samples of pencil thickness (diameter approximately 7 mm) were collected from 32 vines (ID = row-vine number) and tested by commercial Enzyme-linked Immunosorbent Assay (ELISA) kits (Bioreba, Switzerland) for the presence of GLRaV-1, GLRaV-3, GLRaV-4s and GVA according to the manufacturer’s instructions. Viral positive samples were sent to a commercial diagnostics laboratory (Affinity Labs, Adelaide, Australia) for confirmation by multiplex RT-PCR using primers specific to GLRaV-1, GLRaV-3, GLRaV-4s (strains 4, 6 and 9) and GVA. In the second season (2021/2022), samples from asymptomatic GVA (-) vines (n = 8) and from vines with SD symptoms (n=4) were collected. These samples were used to assess viral diversity and phloem transcriptional changes using Illumina next generation sequencing. Lignified canes (diameter approx. 7 mm) were collected from cordon spur positions of each vine following harvest of the 2020/2021 season. The bark was immediately removed with a sharp blade, then phloem tissues were collected and cut into small pieces. The samples were frozen in liquid nitrogen in the field, transported to the University of Adelaide (Waite Campus), then stored at -80 °C until further analysis.

### 3. RNA extraction and sequencing

RNA was extracted from approximately 100 mg of frozen samples according to a modified RNeasy Plant Mini Kit (Qiagen, Germany) procedure (Santi et al., 2013). Samples were homogenized with 1 mL of RLT lysis buffer supplemented with 2.5% (w/v) polyvinylpyrrolidone-40 (PVP-40) and 1% (v/v) β-mercaptoethanol (β -ME). After vortexing, the homogenate was mixed with 1/10 volume of 20% (v/v) N-lauroylsarcosine (Sigma-Aldrich, Australia) and incubated at 70 °C for 10 min. Samples were centrifuged at 13,000 rpm for 5 min and approx. 700 μL of the supernatant was transferred to a QIAshredder column. Contaminated DNA was removed according to On-column DNase digestion protocol (Sigma, USA). The amount of total RNA was quantified using a Qubit fluorometer. Plant ribosomal RNA was depleted using QIAseq FastSelect rRNA Plant Kit (Qiagen, Australia) as described in Liefting et al. (2021). Library preparation and sequencing was performed in the Ramaciotti Centre for Genomics (University of New South Wales, Sydney, Australia). Sequencing libraries were prepared using the Illumina stranded mRNA kit and sequenced with an Illumina NovaSeq 6000 using 2 × 150 bp chemistry on a S4 flow cell.

### 4. Sequence assembly and RNA virus discovery

Illumina paired-end reads were quality trimmed and rRNA depleted using fastp v. 0.23.2 (Chen et al., 2018) and RiboDetector v. 0.2.7 (Deng et al., 2022), respectively. Initial transcriptome assembly was performed using Trinity v. 2.8.5 (Grabherr et al., 2011). Taxonomic classification of assembled contigs was performed using Kaiju v. 1.9.0 against the nr + euk NCBI database. Contigs larger than 3 kb and taxonomically classified as belonging to the Virus superkingdom were retained and queried (blastn) against the GenBank nr nucleotide database for taxonomic assignment. Illumina reads were then mapped back to the transcriptome assembly using Bowtie v. 1.3.1 (Langmead et al., 2009) and the relative abundance of individual contigs were estimated using the RNA-Seq by Expectation Maximization (RSEM) algorithm implemented within Trinity (Li and Dewey, 2011). For phylogenetic analysis, near-complete virus genomes were direction adjusted and aligned using MAFFT v 7.487 (Katoh and Standley, 2013). Maximum likelihood phylogenetic reconstruction was performed using IQ-TREE v. 2.1.2 (Nguyen et al., 2014) and the best substitution model automatically selected using ModelFinder (Kalyaanamoorthy et al., 2017).

### 5. Differential transcript expression analysis

Quality trimmed and rRNA depleted paired-end RNA-Seq reads were used to quantify the transcript abundance in the phloem tissue using annotated CDS sequences of the *Vitis vinifera* reference genome (PN40024.v4) and Kallisto v. 0.48 (Bray et al., 2016). Read count tables were imported into R, features with 0 counts in all samples and a total count < 10 were removed, and differential expression analysis (SD against asymptomatic vines) was performed using the DESeq2 package v.1.24.0 (Love et al., 2014) with default parameters (sample-wise size factor normalization, Cox-Reid dispersion estimate and the Wald test for differential expression) and shrunken Log2 fold-changes. Features with a Log2 fold-change (Log2FC) of |Log2FC| > 1 and an adjusted p-value < 0.005 were considered for further analysis (Table S1). Orthologues to *Arabidopsis thaliana* were obtained from the EnsemblPlants database using the *V. vinifera* (PN40024.v4) gene IDs. Gene ontology enrichment analysis of the ordered differentially expressed transcripts was performed using g:Profiler (Raudvere et al., 2019).

### 6. Sequencing data availability

The sequencing data and viral assembled contigs included in this study are publicly available at NCBI under Bioproject PRJNA1085026.

## Results

### 1. Diversity and abundance of RNA viruses in asymptomatic and SD vines

Four highly symptomatic, GVA (+) vines and eight asymptomatic, GVA (-) vines were sampled for RNA-seq. After quality trimming and *in-silico* rRNA depletion an average of 8.99 million (SD = 0.61 million) reads were obtained per sample (Table S2). Transcript assembly of RNA-seq data allowed for the identification of near-complete genomes of five viral species across four genera of positive-sense ssRNA viruses (*Ampelovirus, Foveavirus, Marafivirus* and *Vitivirus*) that had been previously reported in grapevine (Coetzee et al., 2010) (Figure 1a, Table S3). Genomes matching Grapevine Rupestris Stem Pitting-associated Virus (GRSPaV) and Grapevine Rupestris Vein Feathering Virus (GRVFV) were ubiquitous in the vines regardless of their SD status (Figure 1a). Genomes of GVA and GLRaV-4 were found exclusively in SD-symptomatic samples, with no traces of contigs taxonomically classified as GVA or GLRaV-4 being recovered from any of the asymptomatic samples (Figure 1a). Fragmented contigs corresponding to Grapevine Syrah Virus 1 (GSyV-1) were recovered in three asymptomatic samples (7-9, 14-6, 23-18) (Figure 1a).

**Figure 1.**
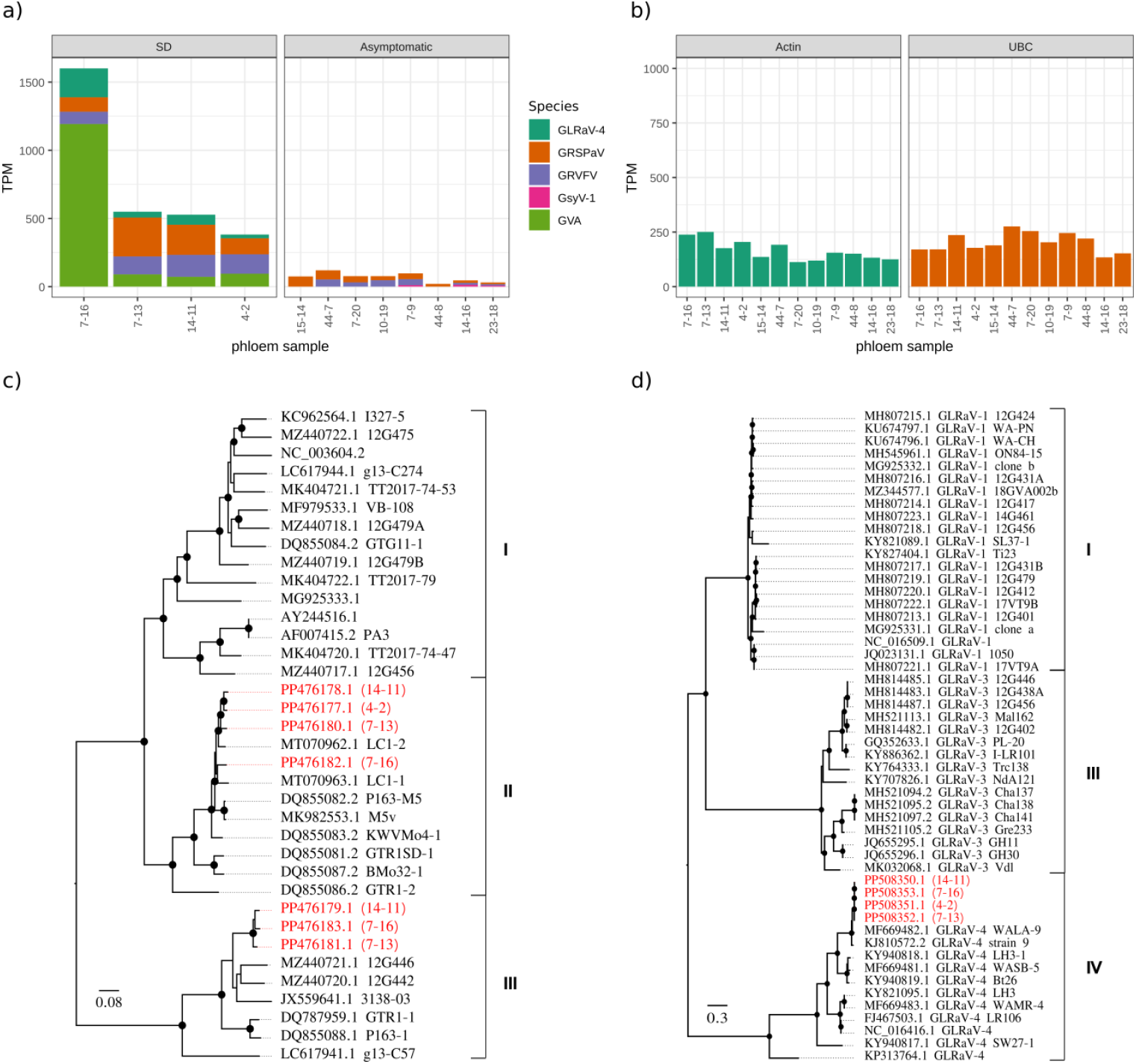
Diversity and abundance of RNA viruses in 12 phloem tissue samples. Transcript relative abundance (TPM, transcripts per million) of viral contigs (a) and two housekeeping genes (b) in samples obtained from asymptomatic and Shiraz Disease (SD) symptomatic grapevines. Maximum likelihood unrooted phylogenetic tree predicted from the sequence alignment of near-complete (c) GVA and (d) *Ampelovirus* GLRaV genomes available at NCBI. Viral genomes assembled in this study are coloured in red, with labels representing the NCBI accession for each assembled genome and the sample ID within brackets. Black circled nodes represent a bootstrap support > 90%.

Virus relative abundance was estimated from the proportion of the total viral reads in each library after removal of rRNA reads. Relative abundance of two housekeeping genes was also estimated, indicating a homogenous transcriptome composition across all samples (Figure 1b). There were large differences in viral abundance between samples, with vines symptomatic for SD displaying more than two-fold increase in total viral abundance relative to asymptomatic vines (Figure 1a). Within SD vines, sample 7-16 displayed the highest total viral abundance as well as the highest abundance of GVA and GLRaV-4 (Figure 1a).

Maximum likelihood phylogenetic reconstruction for GVA and GLRaV-4 was performed using publicly available genomes for GVA (Figure 1c) and GLRaV members of the genus *Ampelovirus* (Figure 1d). Inspection of tree topology confirmed the presence of GVA group II (Figure 1c) and GLRaV-4 strain 9 (Figure 1d) in all SD vines. Symptomatic samples 14-11, 7-16 and 7-13 also contained viral contigs corresponding to GVA group III (Figure 1c).

### 2. Differential gene expression of SD-infected vines

Transcriptomic data from SD and asymptomatic vines were used to investigate regulatory responses in the host grapevine that were associated with viral infection. Transcript expression profiles were compared across phloem tissue samples collected from SD and asymptomatic vines. Unsupervised hierarchical clustering of the normalised transcript counts revealed two distinctive clusters correlating with the SD and asymptomatic samples (Figure S1) and indicated that these biological replicates sampled at identical timepoints, had comparable transcript profiles.

In phloem tissues, a total of 1865 differentially expressed (DE) transcripts were identified after filtering and 1448 and 417 transcripts were up and downregulated, respectively (Table S1). To obtain a global overview of the biological processes affected in SD vines, DE transcripts were annotated using GO terms and then subjected to functional enrichment analysis. The upregulated transcripts were classified into eight GO terms that are primarily involved in phloem development and systemic acquired resistance (SAR) (Figure 2a). For instance, the top upregulated genes were comprised primarily of Sieve Element (SE) Occlusion-like proteins (Figure 2b), which have been linked to SE protein filament formation in eudicotyledons (Anstead et al., 2012; Jekat et al., 2013; Musetti et al., 2013). Several biotic stress responsive genes were also significantly upregulated including Expansin 16 (EXPA16), downy mildew resistant 6 (DMR6), WRKY DNA-binding protein 50, defective in induced resistance 1 (DIR1), ribosomal protein S2 (RPS2) and enhanced disease susceptibility 1 (EDS1) (Figure 2b). Downregulated transcripts were enriched in several GO terms linked to stress response, protein assembly and photosynthesis (Figure 2a). DE analysis also revealed that genes encoding DEAD-box ATP-dependent RNA helicase 46 and heat shock proteins (HSP), as well as photosynthesis-associated genes were significantly downregulated in SD-affected phloem samples at the harvesting stage (Figure 2a, b).

**Figure 2.**
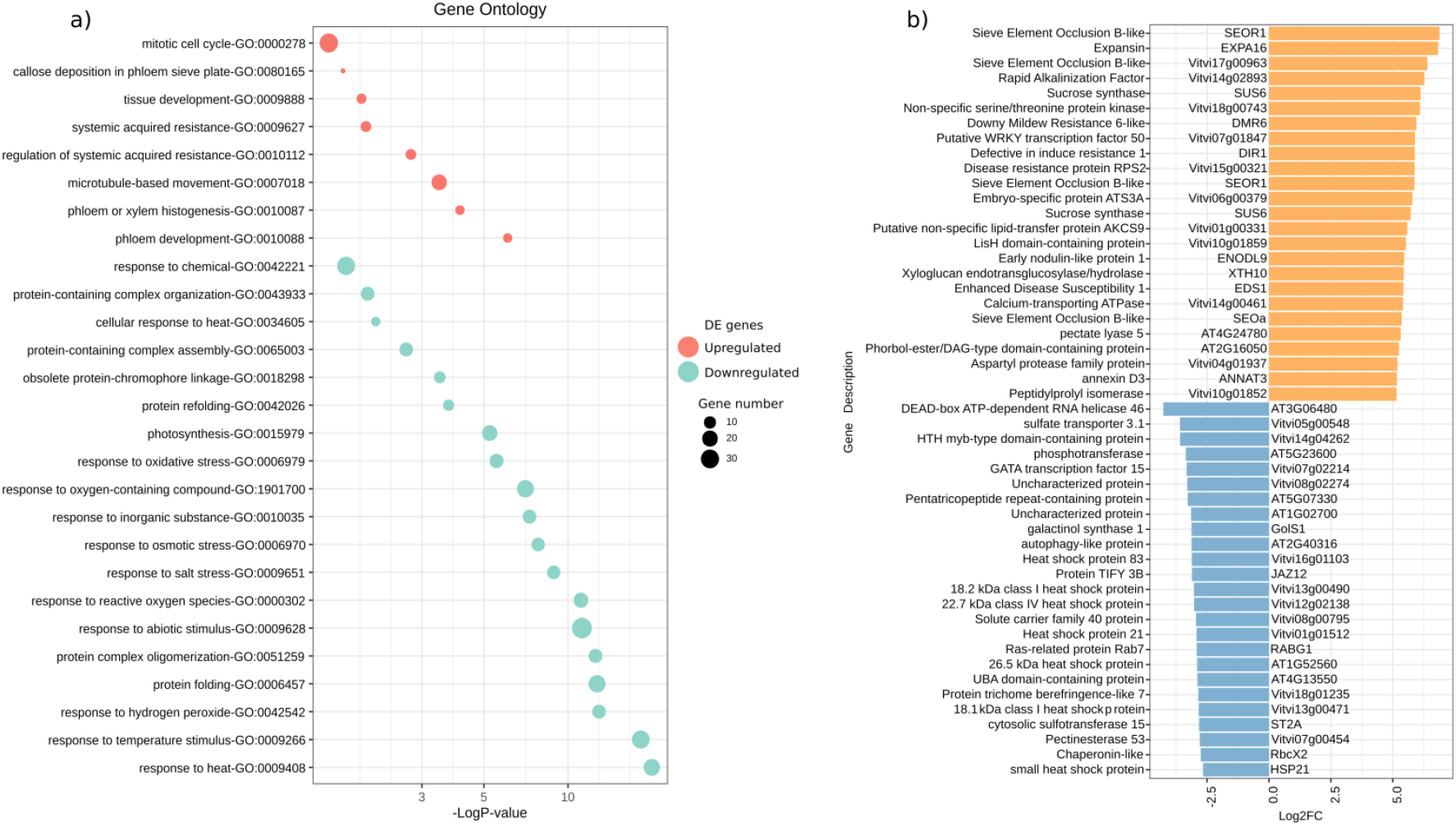
Differential transcript expression analysis between Shiraz Disease (SD) and asymptomatic grapevine samples. a) Gene Ontology (GO) enriched terms in the upregulated and downregulated transcript sets b) Top 50 differentially expressed transcripts between SD and asymptomatic samples. Predicted functional annotations (y axis) and gene names relative to orthologues of *Arabidopsis thaliana* are included when available.

## Discussion

In this study, GVA and GLRaV-4s (strain 9) were predominantly detected in SD-affected vines, whereas GRSPaV and GRVFV were found in both SD-affected and asymptomatic vines. Previous investigations into the viral diversity of SD-symptomatic vines also reported mixed infections involving GVA group II and GLRaV-4 strains 5, 6, and 9 (Wu et al., 2023), suggesting that these two viral species frequently co-exist in SD vines. While we also identified variants of GVA group III in highly symptomatic vines, these variants have previously been reported in vineyards without SD symptoms (Wu et al., 2023).

In mixed infections, beyond virus-host-vector interactions, intra-virus and other complex associations can significantly influence disease establishment and pathogenicity. Co-infections involving multiple viruses have been shown to induce antagonistic, synergistic, or mutually beneficial interactions in several horticultural crops (Mascia and Gallitelli, 2016; Syller and Grupa, 2016). Rowhani et al. (2018) reported that GLRaVs may exhibit synergistic interactions with Vitiviruses, as vines co-infected with GLRaVs (GLRaV-1, 2, or 3) and Vitiviruses (GVA and GVB) showed significantly higher Vitivirus titers, while GLRaV titers remained unchanged compared to control vines without co-infections. Given that the co-occurrence of GVA and GLRaV in vines affected by SD has been documented, further research is needed to elucidate the role of this co-existence in SD pathology.

Investigation of the DE functions within the phloem tissue of SD infected vines provided a clear snapshot of the localised physiological responses of *V. vinifera* to this infection. Similar to the response to other plant pathogens, SD infected vines showed an increased broad-spectrum disease response through upregulation of the systemic acquired resistance (SAR) mechanism, which consisted of increasing the expression of several genes associated with the salicylic acid signalling pathway (Gaffney et al., 1993). *EDS1*, which was amongst the highest upregulated genes, is a positive regulator of salicylic acid biosynthesis and is essential for the Ca2+/calmodulin salicylic-acid-mediated plant immunity (Du et al., 2019). Previous studies on tobacco infected with tobacco mosaic virus have shown that salicylic acid is synthesised at the site of infection and transported to uninfected distal tissues via phloem to establish long-lasting SAR throughout the plant (Shulaev et al., 1995). Salicylic acid enhances the expression of pathogenesis-related (PR) genes that encode proteins which directly affect pathogen growth and proliferation (Zhang et al., 2010). Several salicylic acid-inducible PR genes were also upregulated in SD vines, suggesting that the SAR mechanism might be a defence strategy against SD in *V. vinifera*.

In addition to the upregulation of SAR-associated genes, sieve element occlusion-like genes, calcium transporting ATPase, and genes associated with callose deposition and phloem development were also upregulated in SD-affected phloem tissues. Sieve elements are highly specialised conducting cells that have been designed for long-distance transport of photosynthates, plant growth regulators, and defence and stress signalling molecules. Nutrient-rich phloem tissues serve as an ecological niche for a variety of phloem-borne pathogens and insects and occlusion of phloem sieve plates by callose deposition is a unique and localised defence strategy against phloem-borne viruses to limit their spread throughout the plant (Hao et al., 2008). SE occlusion is dependent on free Ca^2+^ accumulation in the sieve plates (Jiang et al., 2019) and upregulation of calcium transporting ATPase and Expansin 16 (EXPA16), indicates compromised integrity of sieve tubes and suggests that SD-affected vines are attempting to limit the internal spread of the SD viruses by blocking the phloem sieve elements.

Signatures of compromised immunity were also observed within the downregulated functions. Particularly, DEAD-box RNA helicase 46 and several HSPs were downregulated in SD-affected phloem tissues. The DEAD-box proteins are the largest helicase subfamily and their vital roles in plant growth and abiotic/biotic stress tolerance have been reported in several plant species such as Arabidopsis, grapevine, rice, tomato and sweet potato (Jarmoskaite and Russell, 2014; Cai et al., 2018; Wan et al., 2020; Yarra and Xue, 2020; YANG et al., 2022). Ectopic overexpression of rice BIRH1, which encodes a DEAD-box RNA helicase has been shown to enhance expression of oxidative defence genes and disease resistance to *Alternaria brassicicola* and *Pseudomonas syringae* in Arabidopsis (Li et al., 2008). As this evidence supports the putative role of DEAD-box RNA helicases in disease resistance, their downregulation in SD-affected phloem tissues may indicate high disease susceptibility of the symptomatic vines. Furthermore, several studies have demonstrated that HSPs play a major role in the development of innate immunity in plants upon pathogen infections. The plant’s first mode of defence against pathogen infection includes recognition of pathogen-derived pathogen-associated molecular patterns (PAMPs) and effector proteins by pattern recognition receptors (PRRs) and resistance (R) proteins, respectively (Chisholm et al., 2006). HSPs contribute to innate immunity via accumulation of PRRs and stabilisation of R proteins (Hubert et al., 2003) and previous studies have shown that grapevine HSP genes can be upregulated or downregulated in response to virus infections (Espinoza et al., 2007; Gambino et al., 2012; Zhang et al., 2023).

## Conclusion

Collectively, this study shows that vines displaying SD symptoms are infected with multiple viruses, including GVA and GLRaV. The investigation into SD-affected phloem tissues highlighted significant transcriptional changes, including potential roles for the systemic acquired resistance (SAR) mechanism and callose deposition and phloem sieve element occlusion as defence strategies against these viruses. Additionally, the observed downregulation of DEAD-box RNA helicases and heat shock proteins (HSPs) in symptomatic vines suggests a compromised immune response. These findings offer the first insights into the molecular responses of grapevines to SD and provide a foundation for future physiological studies exploring the roles of GLRaV-4, SAR, and phloem occlusion in SD pathology.

## Acknowledgements

This research was made possible through the support of grape growers and winemakers in Australia, via their investment organization, Wine Australia. This was complemented by matching funds provided by the Australian Government. The Australian Wine Research Institute (AWRI) is a part of the Wine Innovation Cluster (WIC) based in Adelaide. We also gratefully acknowledge the financial support provided by the University of Adelaide through the Friendship Fund.

