## Supplementary material for "Viral diversity and phloem transcriptional changes in Grapevine Shiraz disease infected vines": Figure S1

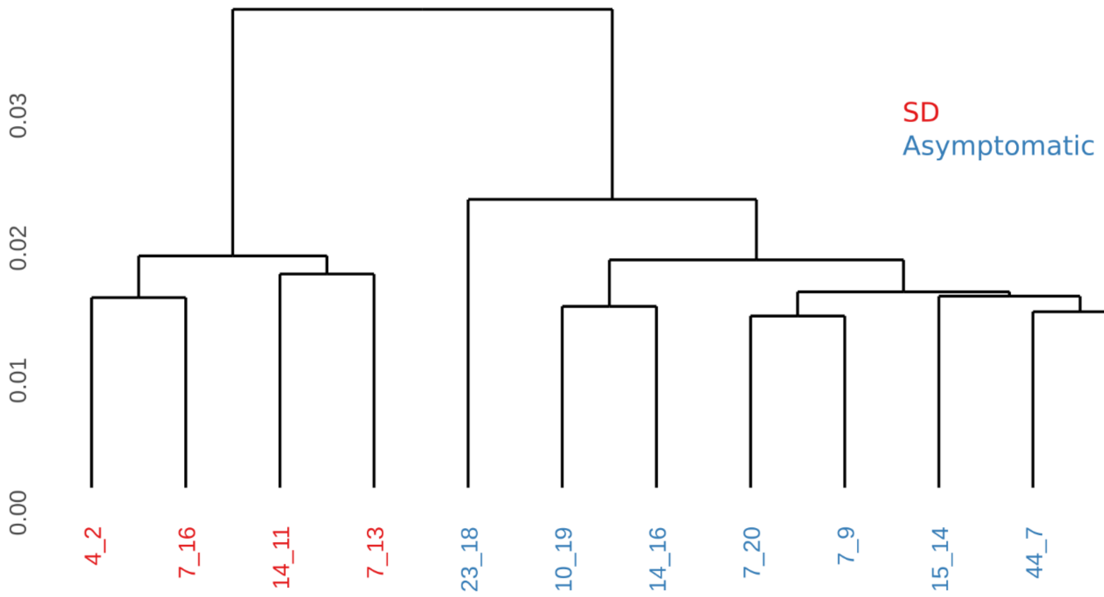


Figure S1. Dendogram of the pearson correlation for the rlog-transformed read counts for the transcriptome of all samples included in the study.
